## Supplementary material for "Elevated Ubiquitin Phosphorylation by PINK1 Contributes to Proteasomal Impairment and Promotes Neurodegeneration": One supplementary table and 7 suplplementary figures

Chen *et al.*

Chun Tang,

**This PDF file includes:**

One supplementary table
Eleven supplementary figures

**Figure 1—table supplementary 1. Clinical and pathological characteristics of brain donors.**

| Donor | NBB/CBB number | Diagnosis | Age at death | Sex | Brain weight (g) | PMD (hrs:min) | Amyloid[2] | Brain region | Neuropathological examination | Source[4] |
|---|---|---|---|---|---|---|---|---|---|---|
| A | 1997-063 | Alzheimer's Disease | 75 | F[1] | 995 | 05:40 | B | CG | | NBB |
| B | 2007-086 | Alzheimer's Disease | 71 | M | 1362 | 05:25 | B | CG | | NBB |
| C | 2019CBB050 | Acute leukemia, secondary malignant bone tumors, renal malignant tumors, pulmonary infection, heart failure | 70 | F | 1272 | 06:20 | O | CG | ND[3] | CBB |
| D | 2022CBB083 | Breast cancer with multiple metastases; acute renal failure; hypertension | 72 | M | 1409 | 13:48 | O | CG | ND | CBB |

1. Abbreviations used, in which F is female, M is male, PMD is postmortem delay, and CG is cingulate gyrus.
2. For amyloid Aβ formation, in which "O" is for absent and "B" is for moderate.
3. ND represents no obvious abnormality detected.
4. Sources of the brain samples are NBB (Netherlands Brain Bank, Netherland) and CBB (National Health and Disease Human Brain Tissue Resource Center, China).

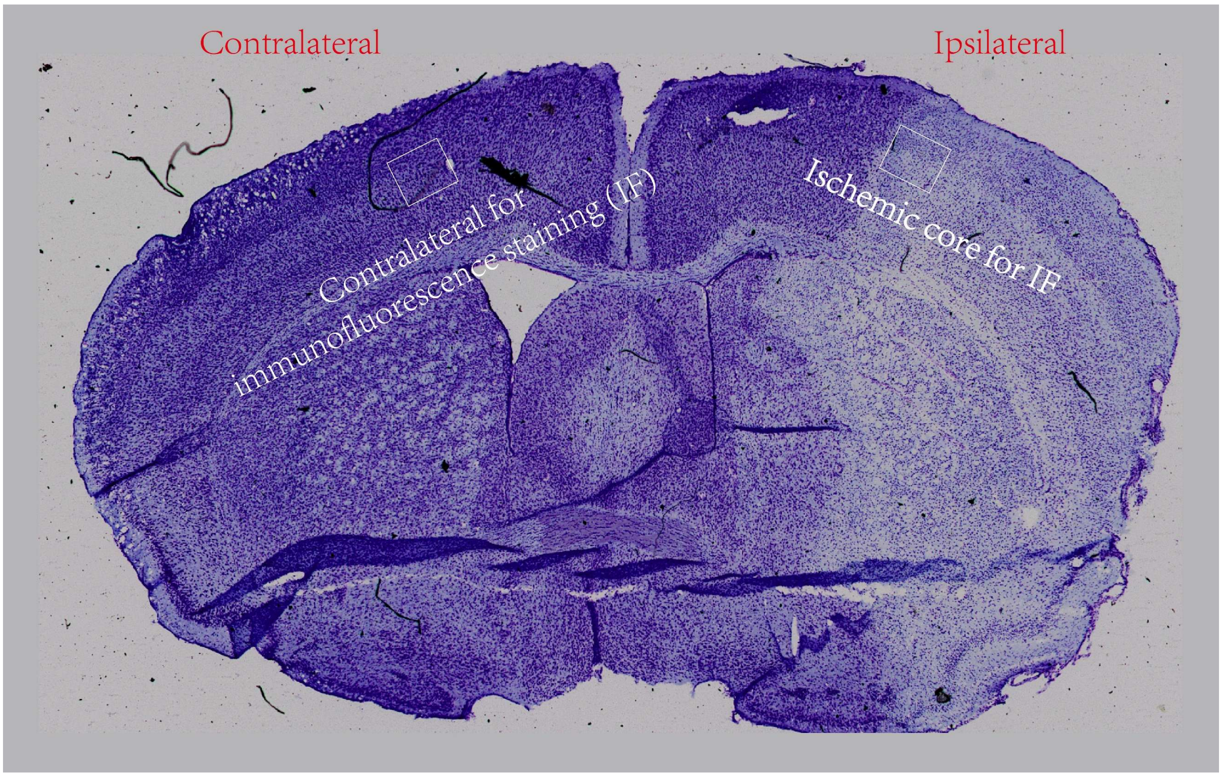


**Figure 1—figure supplementary 1. Nissl staining brain section from a mouse with MCAO to show the brain regions.** The frames show the contralateral and ipsilateral for immunofluorescence staining shown in Figure 1G. The cortex in both sides was separated for Western blot (WB) analysis shown in Figure 4D.

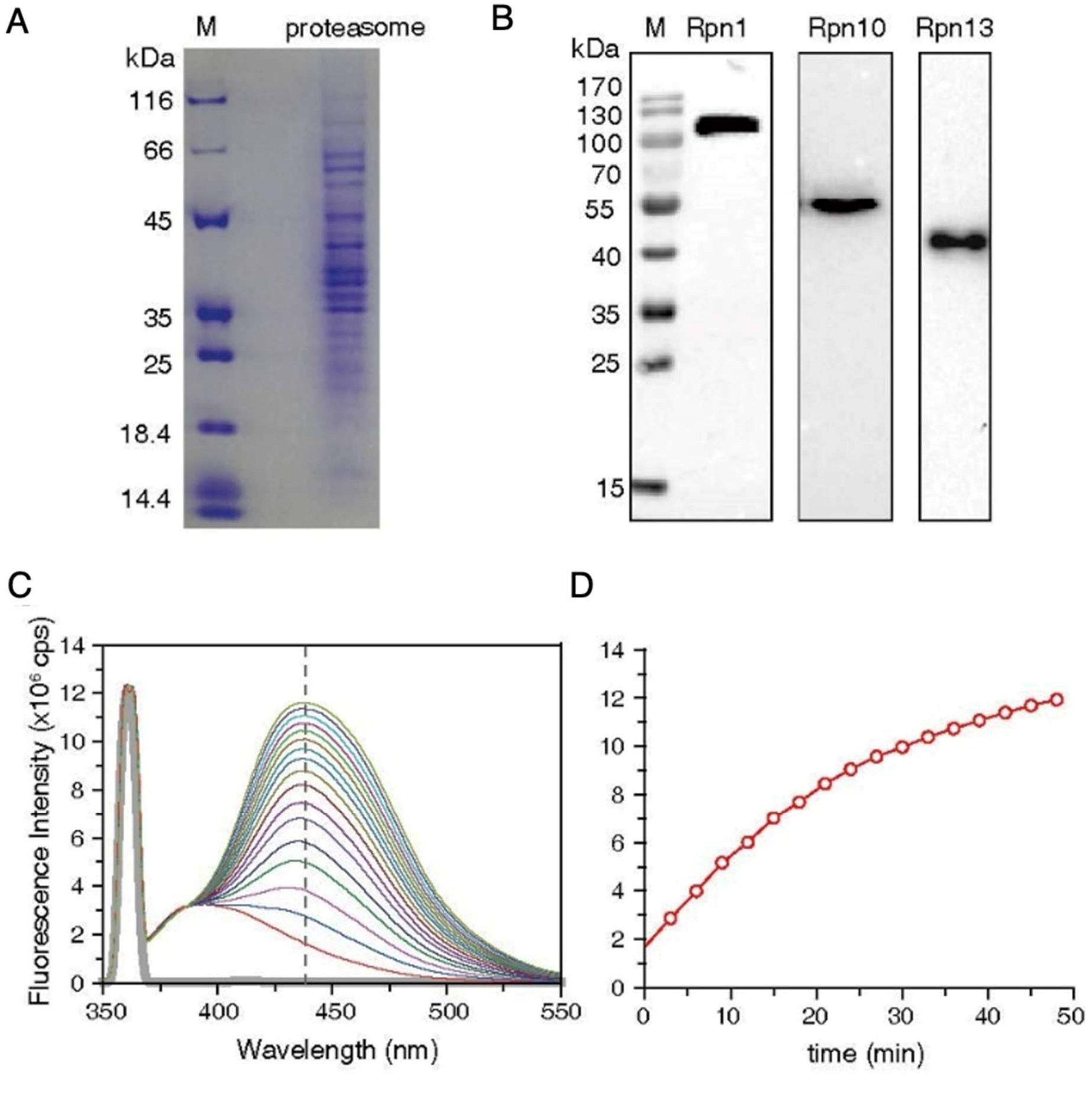


**Figure 3—figure supplementary 1. Preparation of the 26S proteasome for *in vitro* degradation of proteins.** (A) Coomassie blue staining of proteins that make up the proteasome. (B) Western blot identification of the three Ub receptors in the 26S proteasome, including Rpn1, Rpn10, and Rpn13. (C) The determination of *in vitro* proteasome degradation activity using a fluorogenic peptide.

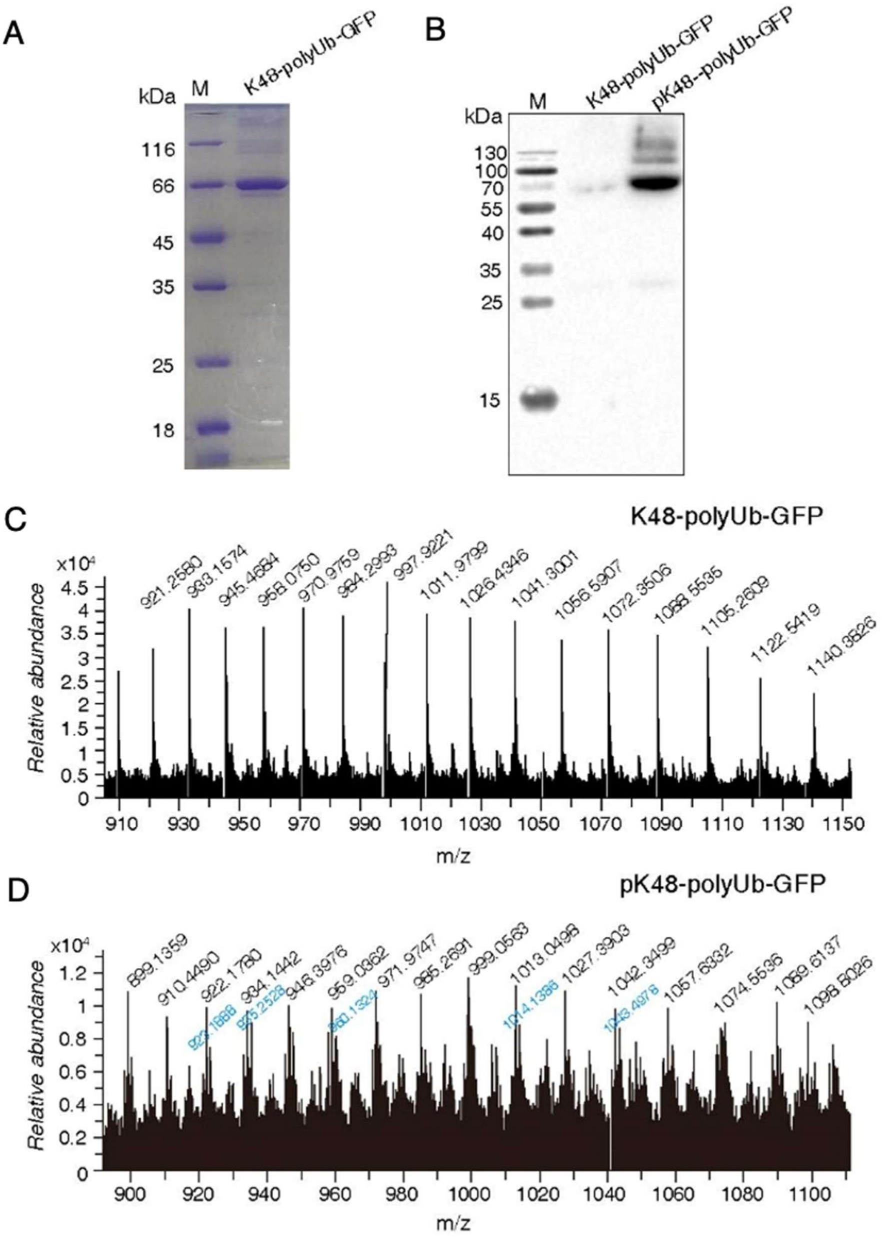


**Figure 3—figure supplementary 2. Preparation of GFP substrate protein modifed with Ub-chain.** The K48-linked tetraubiquitin was covalent attached to Ub-GFP fusion protein through a K48 isopeptide bond through the catalysis of E2-25K. (A) Identification of K48-polyUb-GFP with SDS-PAGE and Coomassie blue staining. (B) Identification of pK48-polyUb-GFP with pUb Western blot. (C and D) ESI-MS analysis of K48-polyUb-GFP and pK48-polyUb-GFP, respectively. The m/z profile indicates that the protein carries 1-2 phosphoryl groups, denoted with black and blue labels.

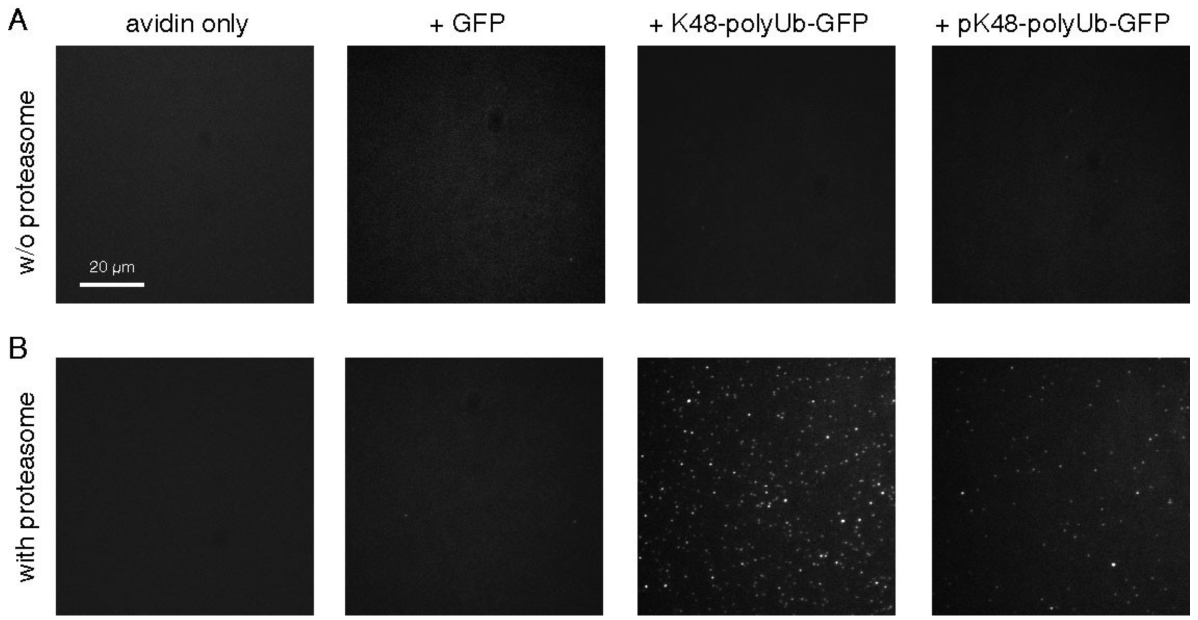


**Figure 3—figure supplementary 3. Visualization of proteasome-associated Ub-modified GFP using TIRF.** (A) Representative TIRF images without the proteasome immobilized on the coverslip. (B) Representative TIRF images with the proteasome immobilized on the coverslip. The immobilized proteasome alone or the application of GFP protein alone could not be visualized. K48-polyUb-GFP can be visualized with TIRF as discrete puncta upon binding to the immobilized proteasome.

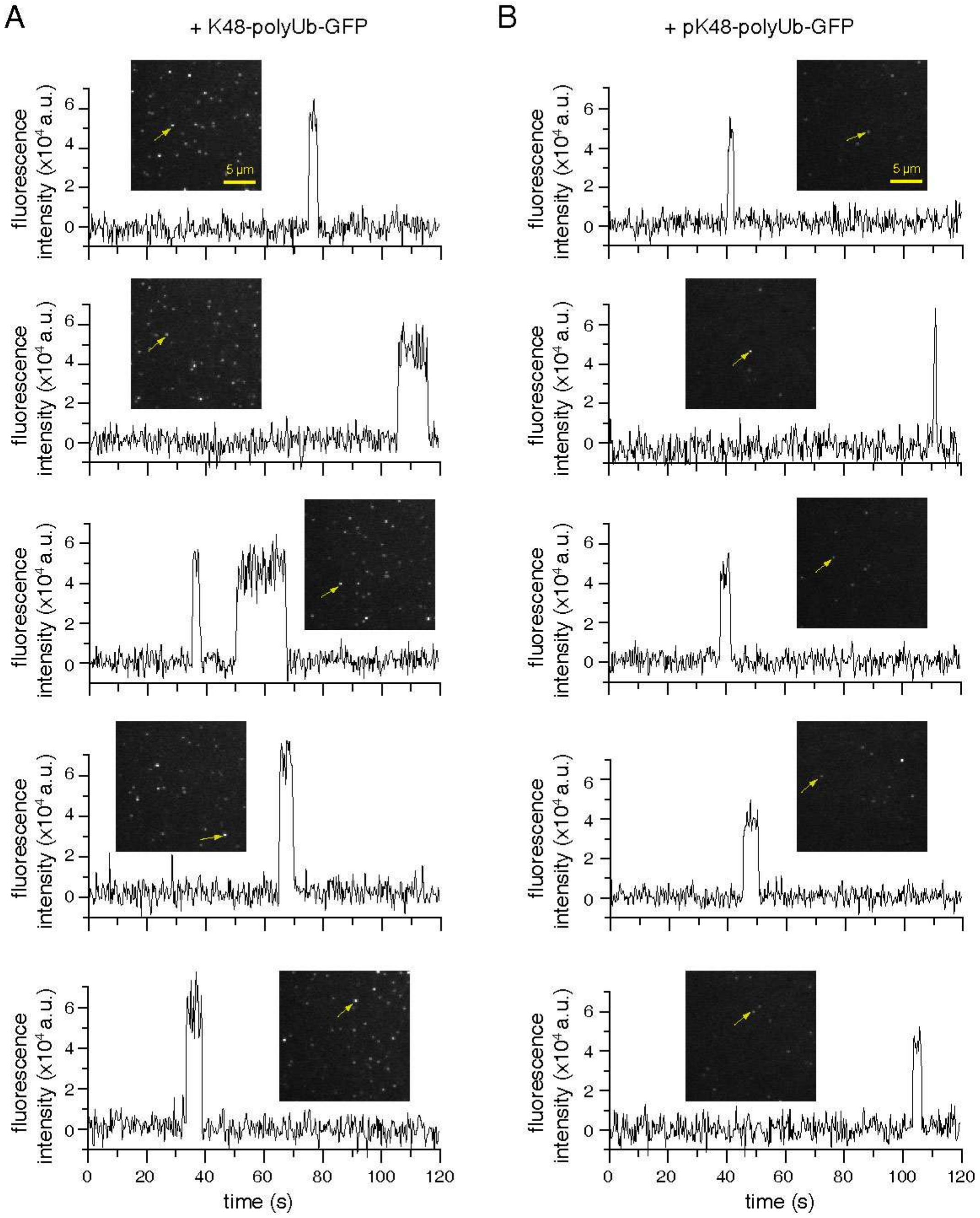


**Figure 3—figure supplementary 4. Time traces of polyUb-tagged GFP substrate visualized with TIRF.** From top to bottom, five representative time traces for K48-polyUb-GFP and pK48-polyUb-GFP, respectively, associated with the immobilized proteasome. The analyzed puncta are indicated with an arrow.

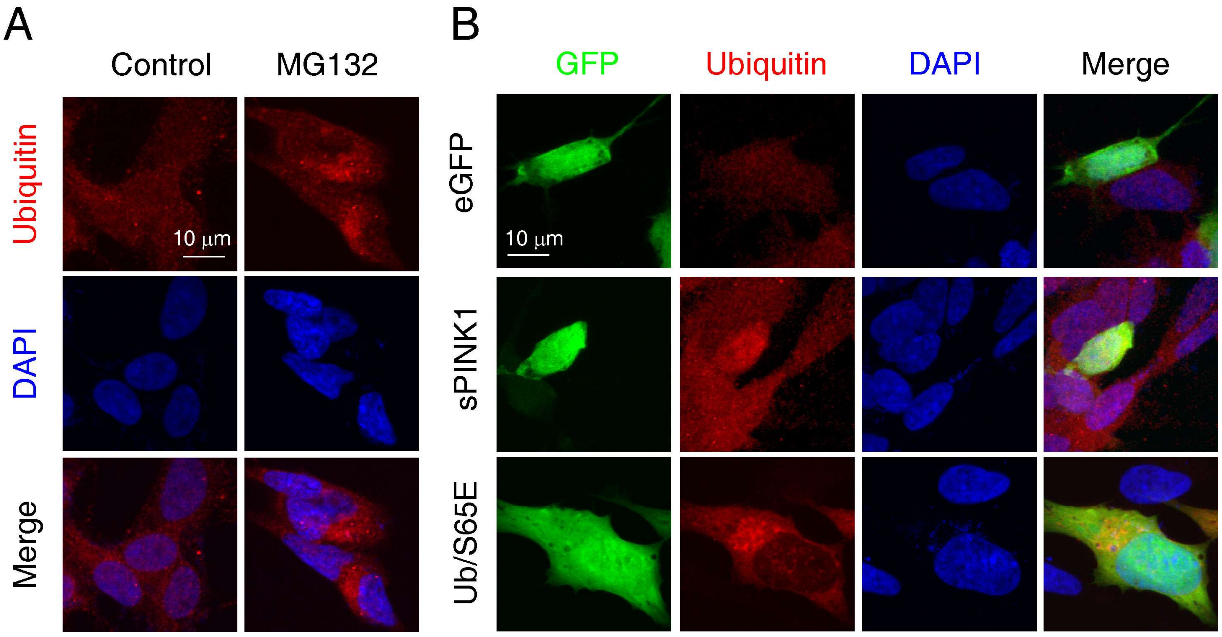


**Figure 5—figure supplementary 1. Over-expression of sPINK1 induced protein aggregation in SH-SY5Y cells**. A. Representative images of immunofluorescent staining of ubiquitin at 24 hours after the treatment of 5 μM MG132. B. Representative images of immunofluorescent staining of ubiquitin at 24 hours after the over-expression of sPINK1 and Ub/S65E.

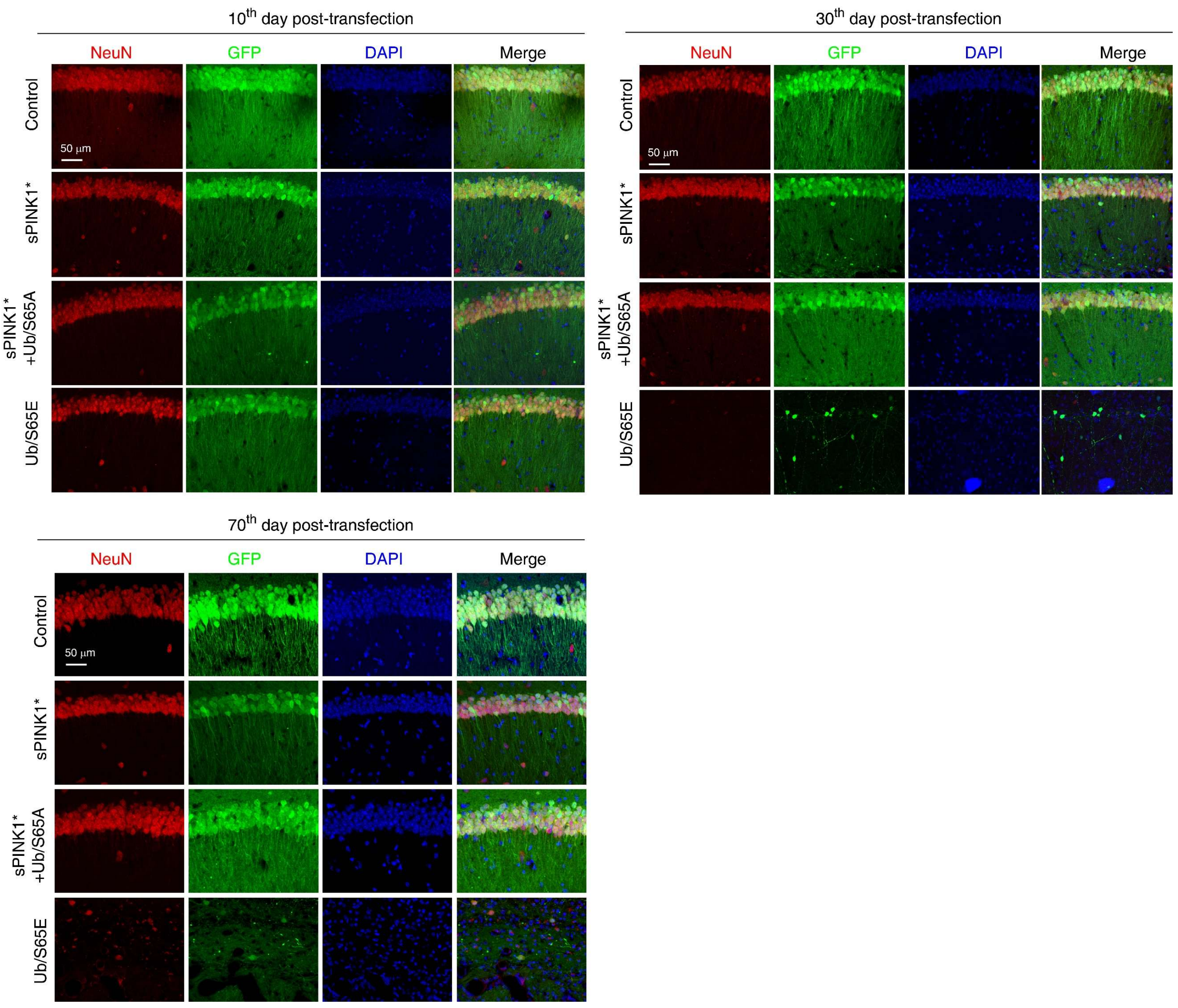


**Figure 5—figure supplementary 2. Representative immunofluorescence images of the hippocampus CA1 regions using anti-NeuN antibody.** Mice were sacrificed at 10-, 30, and 70-days post-transfection. The GFP was introduced to monitor the neurons with AAV transfection.

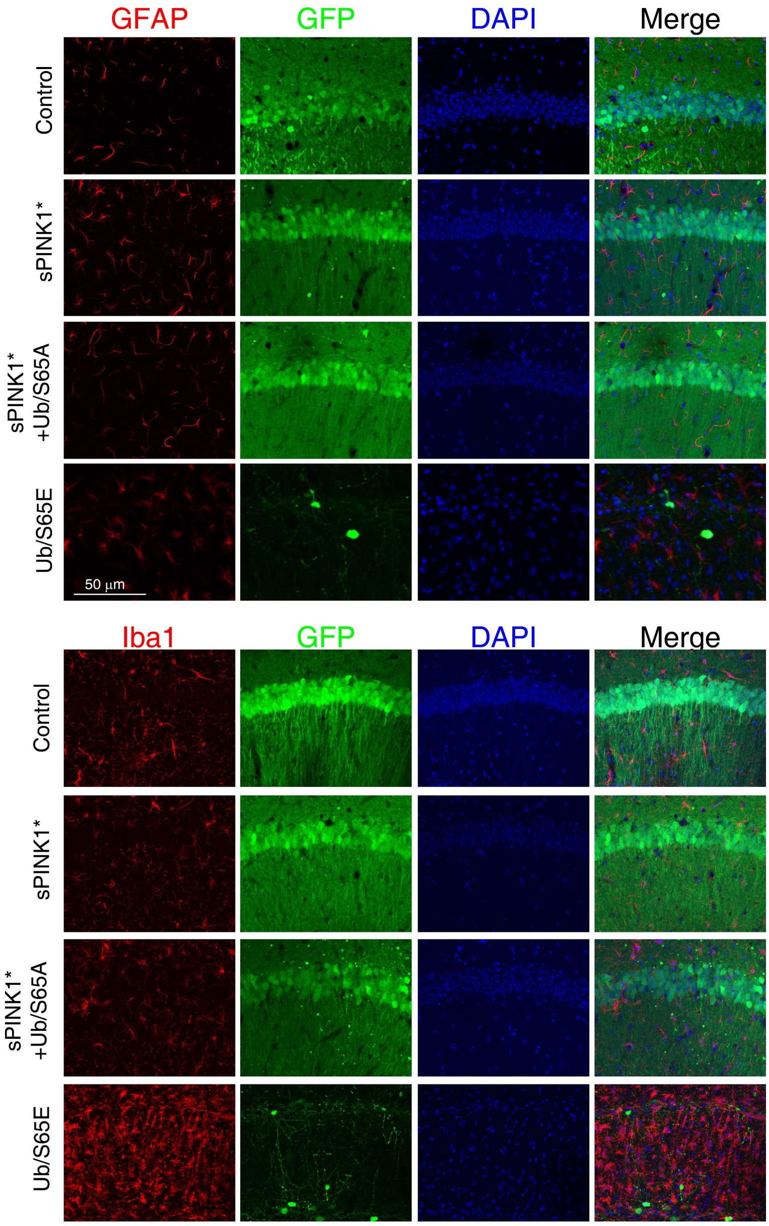


**Figure 5—figure supplementary 3. Representative immunofluorescence images of the hippocampus CA1 regions using anti-GFAP and anti-Iba1 antibody.** Mice were sacrificed at 30-days post-transfection. GFP was introduced to the neurons with AAV transfection.

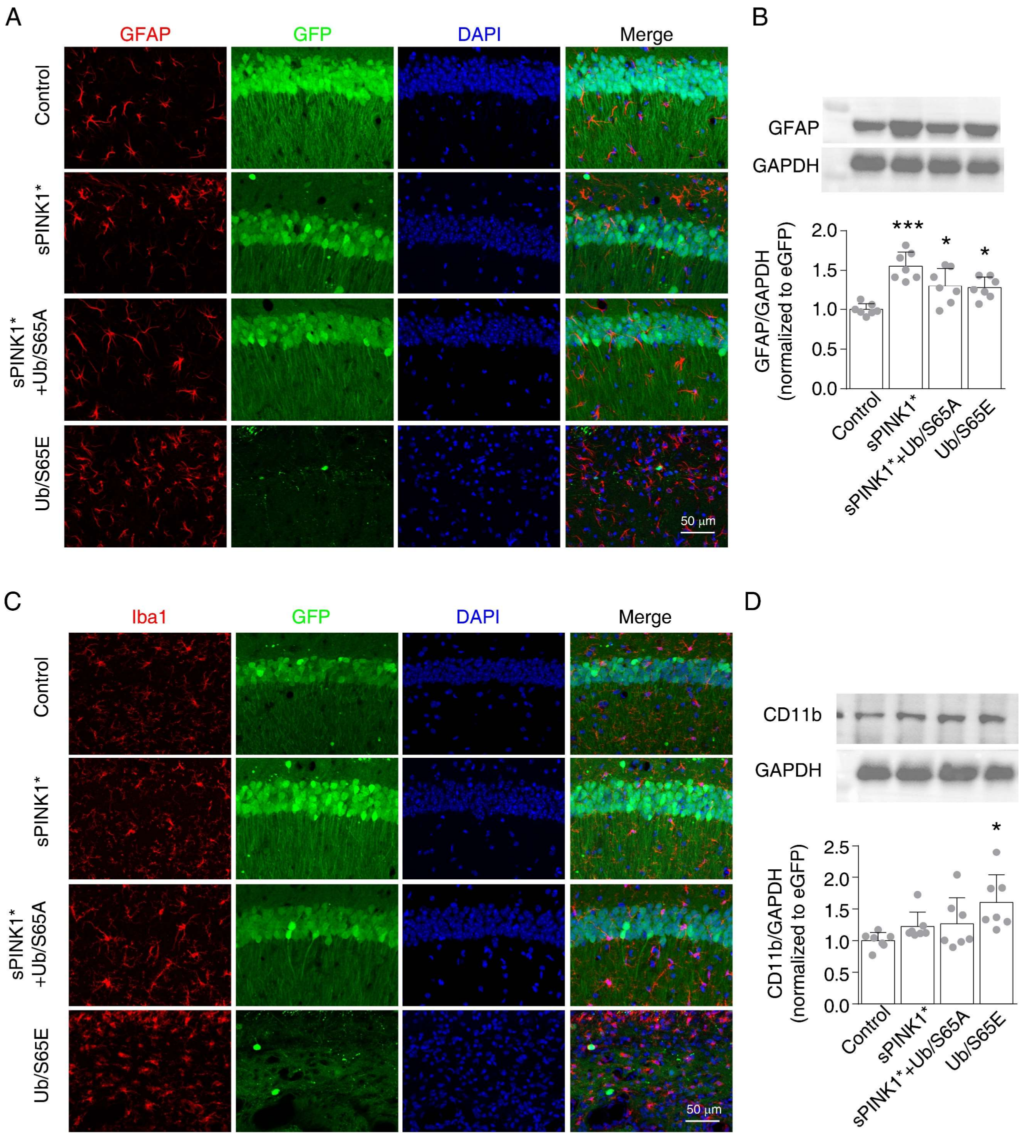


**Figure 5—figure supplementary 4**. **Over-expression of sPINK1* induced gliosis at 70-days post-transfection.** (A) Representative immunofluorescence images of the hippocampus CA1 regions using anti-GFAP antibody. (B) Western blot analysis of GFAP protein. *$P$<0.05, ***$P$<0.001, compared with control, one-way ANOVA. (C) Representative immunofluorescence images of hippocampus CA1 regions immunofluorescent using anti-Iba1 antibody. (D) Western blot analysis of CD11b protein level. *$P$<0.05, compared with control, one-way ANOVA. GFP was introduced to the neurons with AAV transfection.

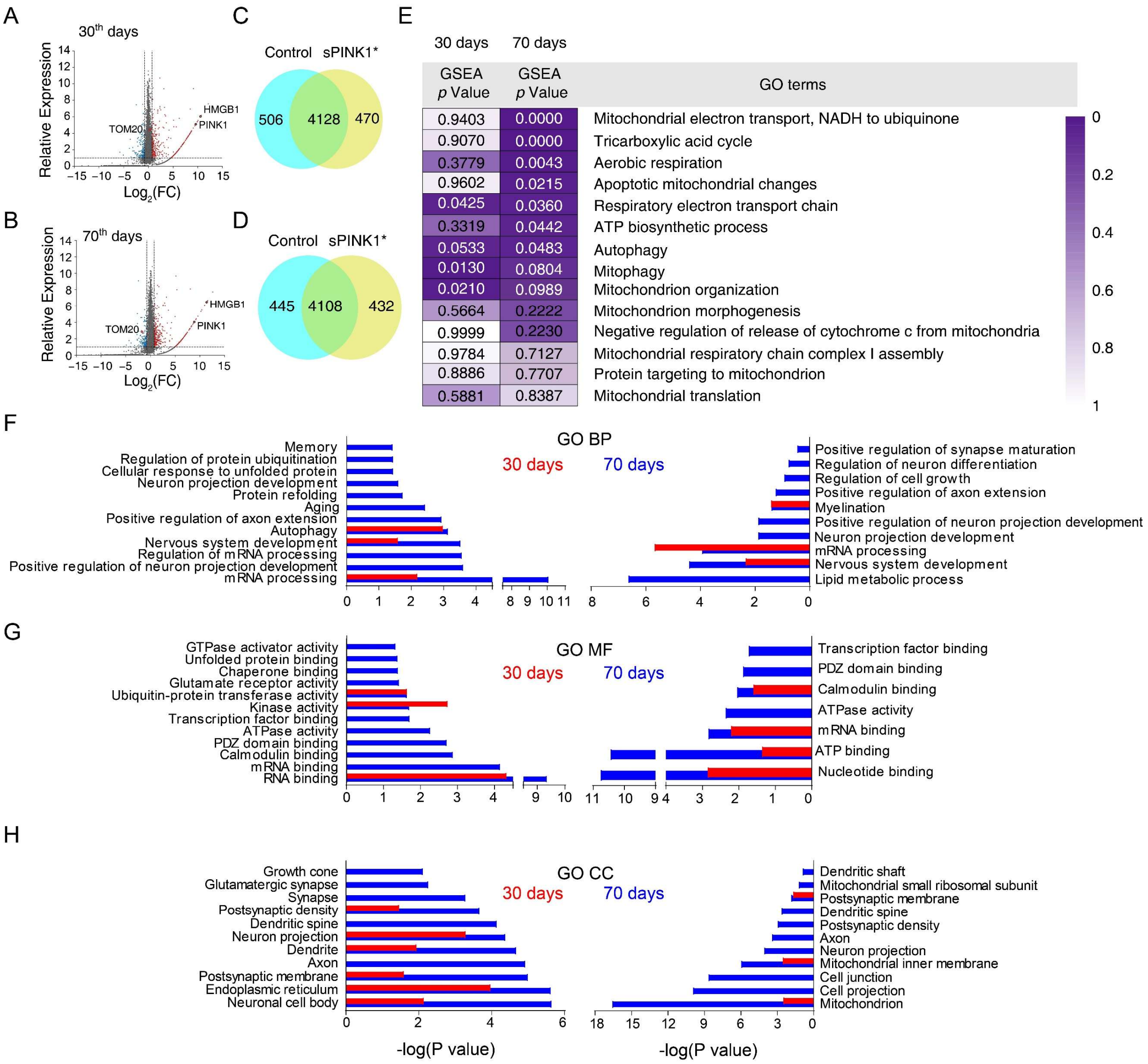


**Figure 5—figure supplementary 5. Proteomics analysis of the mouse hippocampus at 30-and 70-days post-transfection.** (A and B) Volcano plot of the proteomics data from the mouse brains at 30- and 70-days post-transfection, respectively. (C and D) Proteomic analysis revealed differential set of proteins upon sPINK1* over-expression. (E) The GSEA analysis of mitochondria-related GO terms based on the proteomic data, with the statistical significance values color-coded. (F-H) Gene ontology (GO) term analysis of the proteomics data for biological process (BP) (F), molecular function (MF) (G), and cellular component (CC) (H), respectively. Red columns denote the data at 30-days and blue columns at 70-days post-transfection. Left panels denote proteins up-regulated by 2-fold or more, and right panels down-regulated by 50% or more.

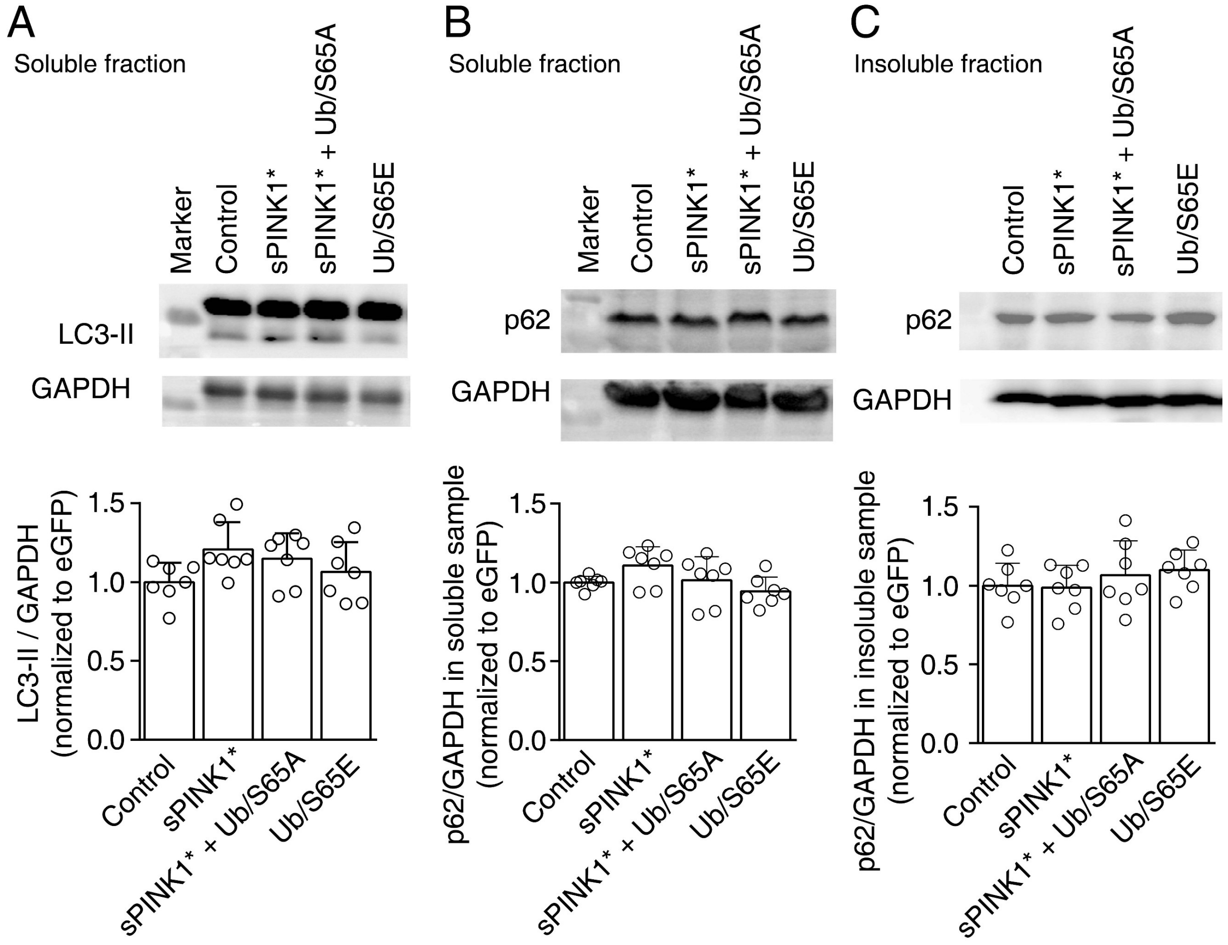


**Figure 5—figure supplementary 6. Elevated pUb level in mouse hippocampus neurons did not perturb the autophagy flow.** Mice were sacrificed on the 70$^{th}$ day post-transfection. The proteins from mouse hippocampus were pooled as soluble samples (A and B) and insoluble sample (C). (A) The level of LC3 in the soluble sample upon AAV transfection of the indicated protein. (B and C) The level of p62 in the soluble (B) and insoluble (C) fractions, respectively. Western blot analysis revealed no statistical differences between the transfections of the different set of proteins.
